# REM sleep reconfigures large-scale network dynamics: a link to its suppressive role in epilepsy

**DOI:** 10.64898/2026.02.27.706763

**Authors:** Gaia Patrone, Maria Giovanna Canu, Gaia Burlando, Monica Roascio, Lorenzo Chiarella, Luca Di Tullio, Laura Tassi, Roberto Mai, Francesco Cardinale, Palva J. Matias, Sheng H. Wang, Maxime O. Baud, Lino Nobili, Gabriele Arnulfo

## Abstract

Converging evidence suggests that human brain activity operates near a critical-like regime in which balanced excitation and inhibition support efficient large-scale communication. The brain’s proximity to criticality may be dynamically reset across the sleep–wake cycle and altered by epilepsy, leading to aberrant oscillatory dynamics.

Building on recent work demonstrating a tripartite interaction between networks synchronization, oscillatory amplitude bistability, and cross-frequency coupling in the human brain that seems to favour epileptic activity, we examined how this interaction, and its underlying large-scale dynamics are modulated across vigilance states.

We analyzed overnight recordings from 20 patients with drug-resistant epilepsy undergoing presurgical evaluation and selected overall 20 minutes of continuous, artifact free stereo-electroencephalography (SEEG) spanning REM sleep, NREM stages N2 and N3, and eyes-closed resting wakefulness. Across states, we quantified phase synchronization, phase–amplitude coupling, bistability and their correlation.

REM sleep was consistently associated with a reduction of these dynamics relative to NREM sleep and wakefulness. Importantly, the canonical correlation between these measures — reflecting the strength of the tripartite interaction — was significantly weaker during REM sleep.

These findings indicate that vigilance states modulate this previously identified multiscale interaction in human brain networks and suggest that the reduced epileptogenicity of REM sleep can be associated with a disruption of coordinated synchronization, coupling, and bistable dynamics at the large-scale network level.

## Introduction

Dynamical-systems theory suggests that cortical networks operate in a critical-like regime characterized by long-range temporal correlations and high sensitivity to perturbations (Hengen and Shew, 2025). This critical state is thought to support flexibility but also makes the network prone to bistability – an irregular switching between low- and high-activity states (Freyer et al., 2009). While a moderate level of bistability within a critical regime appears to support healthy neuronal oscillatory dynamics (Wang et al., 2024, 2023), an excessive or dysregulated bistability may give rise to aberrant oscillatory behaviors.

In this context, large-scale synchronization of neuronal oscillations constitutes as well a key feature of brain functioning, representing an established dynamic for both local excitability regulation (Buzsáki and Draguhn, 2004; Palva and Palva, 2012) and long-range communication including high-frequency oscillations (Arnulfo et al., 2020). Neural oscillations are not stationary but depend on physiological conditions and are continuously reshaped by ongoing changes in neuromodulatory tone and thalamo-cortical drive across the sleep–wake cycle (Nir and de Lecea, 2023).

The probability of epileptiform discharges is also strongly modulated by vigilance states with wakefulness and NREM sleep, particularly N2, generally associated with seizure generation (Eltze et al., 2020; Herman et al., 2001; Minecan et al., 2002; Ng and Pavlova, 2013; Proserpio et al., 2019; Shouse et al., 2000), whereas REM sleep seems to exert a suppressive effect, reducing both interictal discharges and seizure probability (Campana et al., 2017; Frauscher et al., 2016; Frauscher and Gotman, 2019; Moore et al., 2021; Nobili et al., 2025; Shouse et al., 2000). This duality underscores the importance of sleep microstructure in modulating epileptic activity and motivate the investigation of the dynamics behind how specific vigilance states are involved in seizure facilitation or inhibition (Baud et al., 2020). Consistently, in epilepsy, large-scale network dynamics are markedly altered. Evidence indicates that the epileptogenic zone (EZ) typically expresses elevated *β* − *γ* bistability – a novel electrophysiological marker of epileptogenicity (Wang et al., 2024, 2023) – as well as higher levels of within- and cross-frequency coupling (Amiri et al., 2016; Arnulfo et al., 2020). Rather than reflecting isolated phenomena, these signatures point to a multidimensional, interconnected set of processes underlying neuronal communication and varying excitability. Indeed, recent evidence showed that during NREM sleep δ-band phase synchronization, quantified by phase-locking value (PLV), enhances epileptiform discharges, entraining β–γ fast-rhythm bistability within the EZ via phase–amplitude coupling (PAC) (Burlando et al., 2025). Notably, δ-phase activity originating in the non-epileptogenic zone (nEZ) modulated EZ fast activity, especially in N2, supporting a network-level gating of epileptogenic bursts rather than a purely local EZ phenomenon (Burlando et al., 2025). We hypothesize that PAC results both in phase-synchrony in lower and bistability in higher frequency oscillations and constitutes a network-level dynamic that favors epileptic activity. From this hypothesis, we reason that opposing effects of NREM and REM on seizure risk, should be accompanied by pronounced state-dependent differences in these coupling measures, with stronger coupling during NREM and attenuation during REM.

We analyzed continuous overnight stereo-electroencephalography (SEEG) recordings from 20 patients with drug-resistant epilepsy (DRE), undergoing presurgical evaluation. From each subject, we extracted overall 20 minutes of SEEG with four different vigilance states: eyes-closed resting wakefulness (WAKE), NREM stage 2 (N2), NREM stage 3 (N3), and rapid-eye movement (REM).

We investigated whether phase synchronization, phase–amplitude coupling and bistability, as well as their correlation, reflecting a tripartite interaction among these processes, vary across vigilance states and whether REM sleep exerts a specific modulatory effect on these large-scale dynamics. We found a consistent attenuation of these dynamics during REM sleep, potentially linked to the reduced epileptogenicity of REM.

## Methods

### Patients and data acquisition

We analyzed Stereo-EEG data from 20 subjects (age: 29.4 ± 11.2, 14 male) affected by drug resistant epilepsy (DRE) and undergoing pre-surgical clinical assessment for the surgical ablation of the EZ. We acquired monopolar (with shared reference in the white-matter far from the putative epileptogenic zone) local-field potentials (LFPs) from brain tissue with platinum–iridium, multi-lead electrodes. Each electrode has 8 to 18 contacts that were 2 mm long, 0.8 mm thick, and had an inter-contact border-to-border distance of 1.5 mm (DIXI medical, Besancon, France). The anatomical positions and amounts of electrodes varied according to surgical requirements. On average, each subject had 14 ± 2 (mean ± standard deviation) electrodes (range 10–17) with a total of 244 ± 37 electrode contacts (range 180–306, left hemisphere: 242 ± 38, right hemisphere: 223 ± 60 contacts, gray-matter contacts: 112 ± 13).

For each patient, we acquired around 10 minutes of continuous spontaneous brain activity from these patients with eyes closed (resting state) and one overnight recording (7.4 ± 0.9 h) of both SEEG and polysomnography, including EOG, EMG, and scalp EEG electrodes (Fz, Cz, Pz, C3, P3, C4, and P4). We extracted 5 minutes of recording for each of the following vigilance states: resting wakefulness (characterized by uninterrupted spontaneous activity with eyes closed), Clinical experts scored REM and NREM (N2, N3) by visual inspection of polysomnography. Since we aimed to specifically investigate the role of REM sleep in modulating large-scale brain dynamics, we restricted our analyses to 5 minutes of continuous recording per state in order to ensure high-quality, artifact-free data, particularly for REM sleep, which is typically less prevalent and more fragmented.

Recordings included ≥1 EOG channel, chin EMG, and scalp EEG using fronto–central and centro–parietal derivations; scalp lead laterality (right/left) or placement at the vertex was determined individually based on the SEEG investigation and implantation constraints SEEG recordings were acquired with a 192-channel SEEG amplifier system (NIHON-KOHDEN NEUROFAX-1100) at a sampling rate of 1 kHz. Before electrode implantation, the subjects gave written informed consent for participation in research studies and for publication of results pertaining to their data. This study was approved by the ethical committee (ID 939) of the Niguarda Ca’ Granda Hospital, Milan, and was performed according to the Declaration of Helsinki.

### Signal pre-processing

We employed a referencing scheme for SEEG data where contacts in gray matter were referenced to the closest ones in white matter (cWM). This method preserves signal polarity and temporal dynamics while reducing distortions from bipolar referencing and volume conduction artifacts observed in monopolar schemes (Arnulfo et al., 2015). For the following, we refer to SEEG channels or channels to the re-referenced SEEG data.

Prior to the main analysis, line noise at 50 Hz and its harmonics were removed using a bank of notch filters with 1 Hz band-stop width up to 250 Hz. To separate the data into narrow frequency bands we filtered the signals with 40 Morlet wavelets (n° cycles = 7.5) and frequency scales ranging from 2 to 250 Hz, logarithmically spaced.

Epileptic events, such as interictal spikes, exhibit high-amplitude, fast temporal dynamics with widespread spatial diffusion, potentially causing artefacts and artificial increases in synchrony. To address this, before the computation of synchronization metrics, we applied the previously defined wavelet-based time-frequency decomposition, extracting amplitude envelopes per channel and frequency, and segmented signals into 500 ms windows were considered epileptic and rejected if at least 10% of cortical channels concurrently demonstrated amplitude peaks exceeding five standard deviations above their mean amplitude in more than half of the frequencies analyzed.

### Clinical identification of the epileptogenic zone (EZ)

The epileptogenic zone (EZ) was identified through visual inspection of SEEG recordings by experienced epileptologists. Channels were classified as part of the EZ based on the early presence of characteristic peri-ictal and ictal patterns at seizure onset, such as low-voltage fast activity, spike-and-wave complexes, and polyspike-slow wave bursts. For the purposes of this study, both the epileptogenic regions and areas involved in early seizure propagation were grouped together as the epileptogenic zone (EZ). We distinguished them from non-epileptogenic, nearly healthy brain regions (nEZ).

Across the 20 patients, we analyzed a total of 1575 nEZ channels (mean per patient: 78.75 ± 17.28) and 649 EZ channels (mean per patient: 32.45 ± 13.97).

Seventeen of 20 participants were operated, three out of the 20 subjects had no surgery. Post-surgical evaluations confirmed the correct localization of the EZ in the 65% of the operated cohort (9 *Engel* IA, 2 *Engel* IIA, 4 *Engel* IIIA, 2 *Engel* IVA).

### Phase synchronization

We estimated phase synchronization between channels at the individual subject level using the Phase Locking Value (PLV) (Aydore et al., 2013). This measure quantifies the consistency of phase differences between signals, providing an estimate of instantaneous phase synchronization. PLV values range from 0 to 1, with values approaching 1 indicating stronger synchronization, meaning the phase difference between two signals remains consistent over time.

First, we performed a time-frequency decomposition by filtering the data using Morlet-wavelets (see Methods, Signal pre-processing). Subsequently, PLV was computed as:

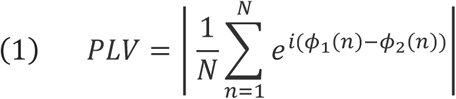

where *ϕ*_1_(*n*) and *ϕ*_2_(*n*) are the instantaneous phases time series from two recording channels, and N is the number of time points.

### Cross-frequency coupling

Two time series could be defined as cross-frequency phase-amplitude coupled if the amplitude fluctuations of fast rhythms the phase of the slow oscillations. We evaluated cross-frequency interactions at the individual subject level using Phase-Amplitude Coupling (PAC) (Siebenhühner et al., 2020). PAC quantifies the extent to which the phase of slower oscillations relates the amplitude of faster oscillations within or across SEEG channels, where higher PAC values indicate greater modulation. In this study, we estimated PAC with the PLV defined as:

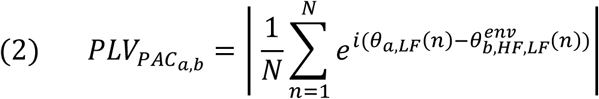

where *θ*_*a*,*LF*_(*n*) is the phase of the signal filtered with a Morlet filter at LF and *θ*_*b*,*HF*,*LF*_(*n*) is the phase of the amplitude envelope of the HF signal filtered with a Morlet filter at LF. Local PAC was obtained where *a* = *b*, interareal PAC where *a* ≠ *b*.

To estimate Phase-Amplitude Coupling (PAC), we constructed a design matrix using ten low-frequency (LF) components, spanning from 1.2 to 68.1 Hz, and 19 frequency ratios ranging from 1:2 to 1:100 (slow : fast). After computing PAC matrices in this LF-to-ratio format, we converted them into a more interpretable low-frequency-to-high-frequency representation and applied interpolation to fill in any missing values. The resulting values m

### Network bistability

Neuronal bistability refers to the irregular alternation between an up and a down state in an oscillatory signal. This phenomenon, which can be driven by positive feedback mechanisms, is theoretically associated with a first-order phase transition (Cowan et al., 2016; Freyer et al., 2012). To quantify bistability, we computed the bistability index (BiS) by fitting the probability distribution of the narrow-band power time series of each sleep segment with both single-(Eq. 3) and bi-exponential models (Eq. 4), selecting the best-fitting model based on the Bayesian Information Criterion (BIC). The single-exponential model is defined as follows:

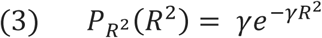

where *γ* is the exponent, and the bi-exponential model is defined as:

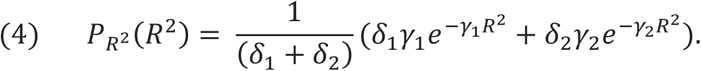

where *γ*_1_, *γ*_2_ are the two exponents and *δ*_1_, *δ*_2_ are weighting factors. The BIC was computed for the single- and bi-exponential fitting as follows:

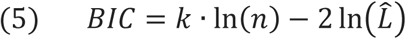

where *n* is the number of samples, 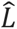 is the likelihood function, *k* is the number of free parameters in the model: *k* = 1 for single-exponential *BIC*_*Exp*_ and *k* = 4 for bi-exponential model *BIC*_*biE*_. A better-fitted model corresponds to a smaller BIC value.

The difference in BIC between single- and bi-exponential is obtained as:

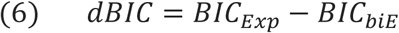

and the bistability index BiS as:

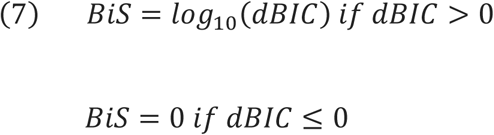

A BiS near zero suggests that the time series is better described by a single-exponential distribution, whereas when values above 3 the bi-exponential model more accurately captures the distribution.

### Canonical Correlation Analysis

Canonical Correlation Analysis (CCA) is a multivariate statistical method used to identify the strongest associations between two sets of variables. It has been widely applied in the analysis of high-dimensional data in neuroscience. This technique aims to find canonical weight vectors, u and v, which maximize the correlations between projected data:

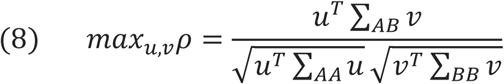

where *ΣAA* = (*A*_*T*_ *A*)/*n* is the covariance matrix for the data array *A* (*e*.*g*., inward-PAC, size *n* × *a, n* is channels number) and *ΣBB* = (*B*_*T*_*B*)/*n* is the covariance matrix for data array *B* (*e*.*g*., bistability, size *n* × *b*). *ΣAB* = (*A*_*T*_ *B*)/*n* is the cross-covariance of *A* and *B*. Solving the generalized eigenvalue problem:

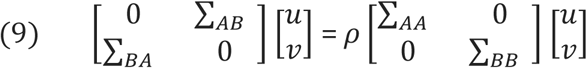

yields the canonical correlation coefficients *ρ* and the weight vectors *u* and *v*. The canonical variates (CV) are computed as:

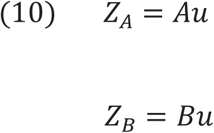

where *ZA* and *ZB* (both with size *n* × 1) are the CVs of *A* and *B* ; *u* (*a* × 1) and *v* (*b* × 1) are the first canonical weight vectors.

We used CCA to explore the relationships between phase dynamics, exploiting outward-PAC (how the amplitude of a faster oscillation is modulated by the phases of slower waves) and phase synchrony, and between amplitude-based metrics inward-PAC (how the phase of a slower wave modulates the amplitudes of faster activity) and bistability.

Inward- and outward-PAC vectors were obtained by averaging each PAC connectivity matrix column-wise and row-wise, respectively. Here, ‘inward’ and ‘outward’ describe the organization of the PAC connectivity graph and should not be interpreted as implying directionality or causality between the slow and fast oscillations. The inward vector summarizes, for each channel, the extent to which its amplitude is statistically coupled to the phase of other channels. Conversely, the outward vector summarizes, for each channel, the extent to which its phase is statistically coupled with the amplitudes of other channels. The CCA was carried out using the Python scikit-learn module.

To quantify how strongly a single channel is phase-synchronized with the specific parts of the network we computed a selective Node Strength (SNS) for each selected channel as the average Phase Locking Value (PLV) between that channel and all the others of the considered zone (EZ, nEZ, nEZ-EZ).

All the metrics where calculated exploiting the Python toolbox “CROCOpy” for the evaluation of brain criticality and connectivity (Myrov et al., 2026).

### Surrogates’ construction and statistical analyses

To recreate the null-hypothesis distributions for both PLV and PAC, we created a set surrogate data using a time-rotation approach. For one time-series in each pairwise comparisons, we randomly selected a time point (k) to split the narrow-band time series into two segments:

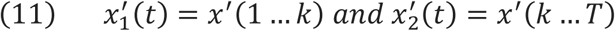

These were then reordered and concatenated in reverse to create a surrogate time series, *x*′*surr*(*t*) = [*x*_2_′, *x*_1_′]. Surrogate PLV and surrogate PAC values were then computed for all channel pairs.

A preliminary analysis to assess for significant differences across all vigilance states was computed by Friedman test, while effect sizes were computed by Kendall’s W.

Post-hoc paired comparisons were assessed using the Wilcoxon signed-rank test. For PAC, the test was applied to each low-frequency–high-frequency (LF–HF) pairs. In the case of BiS and PLV, the test was performed at each selected frequency.

Effect sizes for between-group differences were estimated using Cohen’s d, while confidence intervals under the null hypothesis were derived from 10,000 label-shuffled surrogate datasets.

To address the issue of multiple comparisons, p-values were corrected using the Benjamini-Hochberg (BH) procedure, with statistical significance defined as *p* < 0.05 after correction.

Differences in canonical correlation coefficients (REM vs. N2, N3, WAKE) were instead tested against a null distribution generated through permutation of Y matrices across states (10,000 iterations). Statistical significance was evaluated by comparing observed differences with the permutation distribution, and p-values were corrected for multiple comparisons by using the Bonferroni method.

## Results

Our cohort included SEEG from 20 patients with heterogeneous types of drug-resistant epilepsy (9 temporal lobe epilepsy; 11 extra-temporal lobe epilepsy) undergoing presurgical evaluation (29.4 ± 11.2 years, 14 male). For each subject, we examined 5 min recording for each vigilance states: eyes-closed resting wakefulness, REM, N2, and N3.

Channel pairs were grouped into four channel-pair configurations: nEZ → nEZ, nEZ → EZ, EZ → nEZ, and EZ → EZ (Fig. 1a). Because PLV is a symmetric measure, phase synchronization analyses collapsed cross-region pairs into a single nEZ–EZ group, yielding three categories. In contrast, bistability is a univariate metric that yields a single value per channel; therefore, bistability analyses distinguished only between EZ and nEZ channels rather than channel pairs.

**Fig 1:**
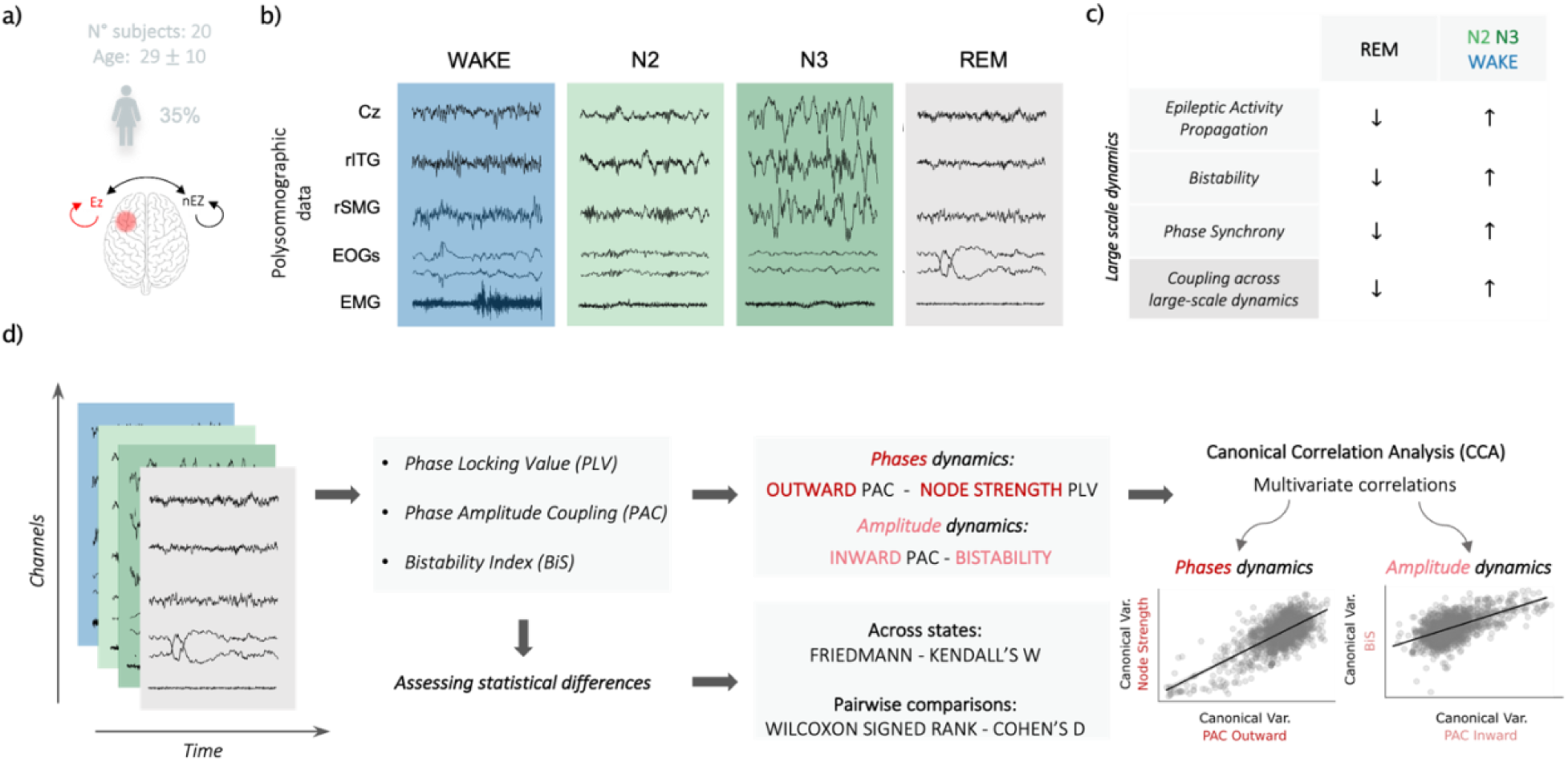
Schematic of the study. a) Above, demographic characteristics of the study cohort, including n° of subjects, sex distribution and age (mean ± SD), below, the definition of the four channel-pair configurations used for the analysis, based on the epileptogenicity of the electrode channels (EZ, epileptogenic zone; nEZ, non-epileptogenic zone): nEZ → nEZ, EZ → nEZ, nEZ → EZ, EZ → EZ; b) Example of polysomnographic data from the analyzed sleep stages: resting state with eyes-closed (WAKE), N2, N3 and REM; c) Hypothesis: not only epileptic activity propagation, network instability and synchrony but also their coupling are reduced in REM compared to the other vigilance states; d) Pipeline of the study: SEEG signals were used to compute phase synchronization (PLV), phase–amplitude coupling (PAC), and bistability (BiS). State-dependent differences were assessed using Friedman tests (Kendall’s W) and Wilcoxon signed-rank post hoc comparisons (Cohen’s d). Phase dynamics were characterized by outward PAC and PLV-derived node strength, whereas amplitude dynamics included inward PAC and bistability. Finally, canonical correlation analysis (CCA) was applied to examine multivariate associations between phase- and amplitude-related dynamics across vigilance states.

### Reduced θ − α phase synchrony during REM sleep

Compared to all other states, REM sleep exhibited a distinct oscillatory coupling profile.

Specifically, REM lacked the prominent α-band synchronization observed in the other states, as well as the θ and spindle-range coupling characteristic of N2 and N3 sleep (Wilcoxon signed-rank test, BH-corrected, p < 0.05; Fig. 2b, Suppl. Fig. 2). These reductions in the θ–α range (4-13 Hz) reached up to ∼30%, and were associated with large effect sizes in nEZ–nEZ and nEZ–EZ pairs (|d| > 0.8, Suppl. Fig. 2), whereas effects in EZ–EZ pairs were smaller and largely confined to α frequencies (0.2 < |d| < 0.5, Suppl. Fig. 2).

**Fig 2:**
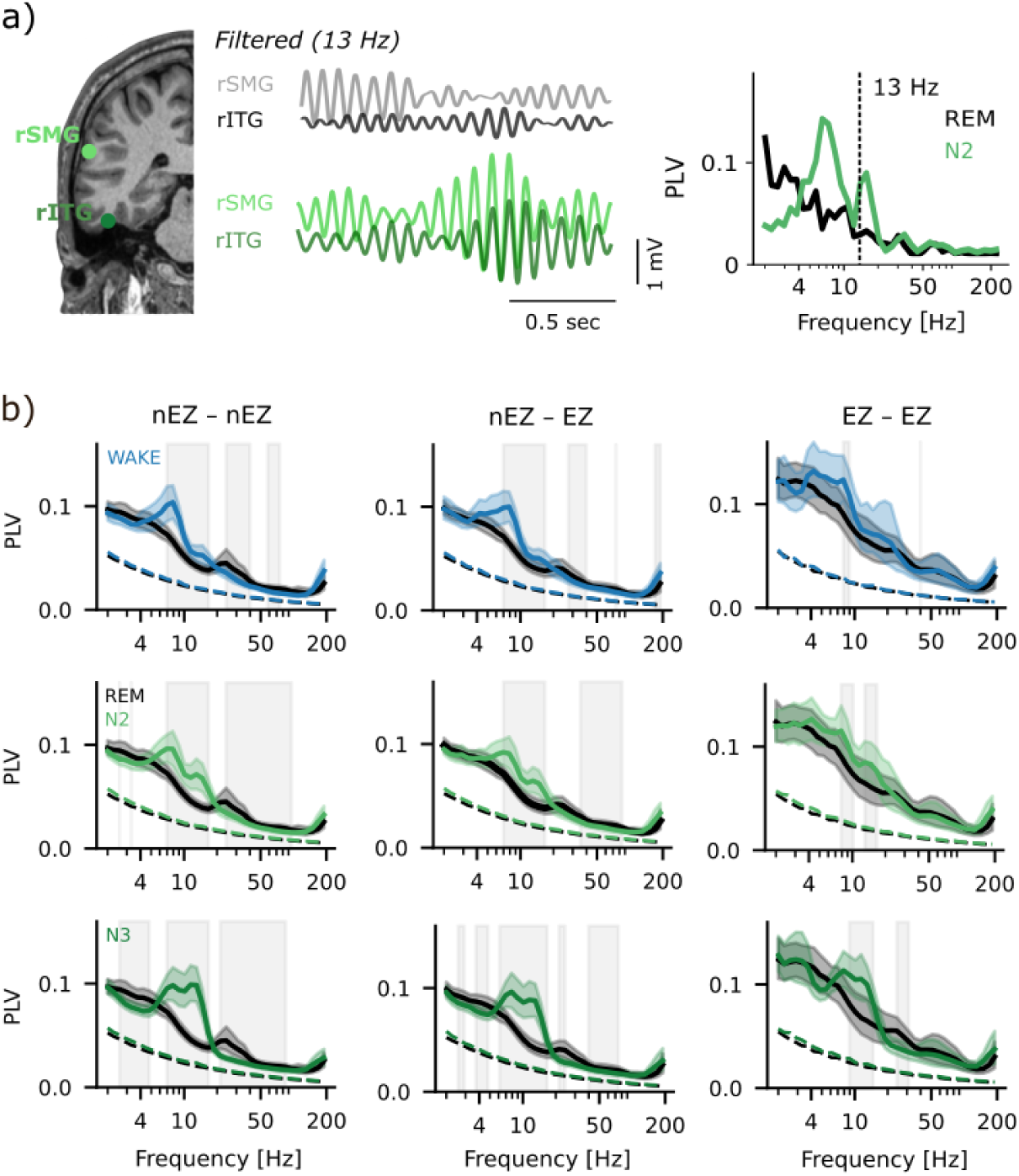
Reduced θ − α phase synchrony during REM sleep. Phase-Locking Value (PLV) computation between a single channel pair (channel 1: right inferior temporal gyrus (rIGT), channel 2: right supramarginal gyrus (rSMG)) for REM and N2; a) Comparison between PLVs spectral profiles across all states showing p-values (Friedman test, BH-corrected; grey lines) and effect sizes (Kendall’s W; red lines), the grey dashed line represent the significance level (*α*). b) Spectral profiles for phase locking values for nEZ–nEZ, nEZ–EZ, EZ–EZ channel pairs, showing mean PLV spectra (solid lines), with corresponding surrogates (dashed lines). Colored shaded areas indicate bootstrapped, two-tailed 95% confidence intervals (N=1000). Grey vertical bands highlight frequency ranges with significant differences (*p* < 0.05) in PLV (Wilcoxon signed-rank test, BH-corrected)z

In contrast, REM uniquely displayed enhanced synchronization in the β–low-γ range (20–100 Hz), with PLV values up to 35% higher than N3 (p < 0.05, |d| > 0.8, Fig. 2b, Suppl. Fig. 2), and moderately increased relative to N2 and wake (0.5 < |d| < 0.8, Fig. 2b, Suppl. Fig. 2), primarily in nEZ–nEZ and nEZ–EZ pairs. Additionally, REM showed a modest increase in δ–low-θ (2–5 Hz) synchronization compared to N3 in the same channel groups (p < 0.05, 0.5 < |d| < 0.8, Suppl. Fig. 2).

Overall, REM was characterized by a redistribution of phase synchronization across frequencies: reduced low-frequency and α coupling typical of wake and NREM sleep, alongside a relative enhancement of higher-frequency synchronization, particularly in networks involving non-epileptogenic tissue.

### Reduced cross-frequency coupling during REM

To investigate state-dependent cross-frequency modulation of neural oscillations, we compared normalized phase–amplitude coupling (nPAC, normalized by dividing for the chance level estimated as the mean of 100 surrogates, see Methods), between REM sleep and the other vigilance states, which assess whether local faster oscillations were correlated to the phase of distant slower oscillations.

A small degree of PAC is present during REM sleep (Amiri et al., 2016; von Ellenrieder et al., 2020), yet in our data REM consistently exhibited a significant reduction of nPAC over a wide range of frequency pairs (*p* < 0.05, Wilcoxon signed-rank test, BH-corrected; Fig. 3b, Suppl. Fig. 1). This decrease was observed across all four channel-pair configurations and in comparisons with each of the other vigilance states.

**Fig 3:**
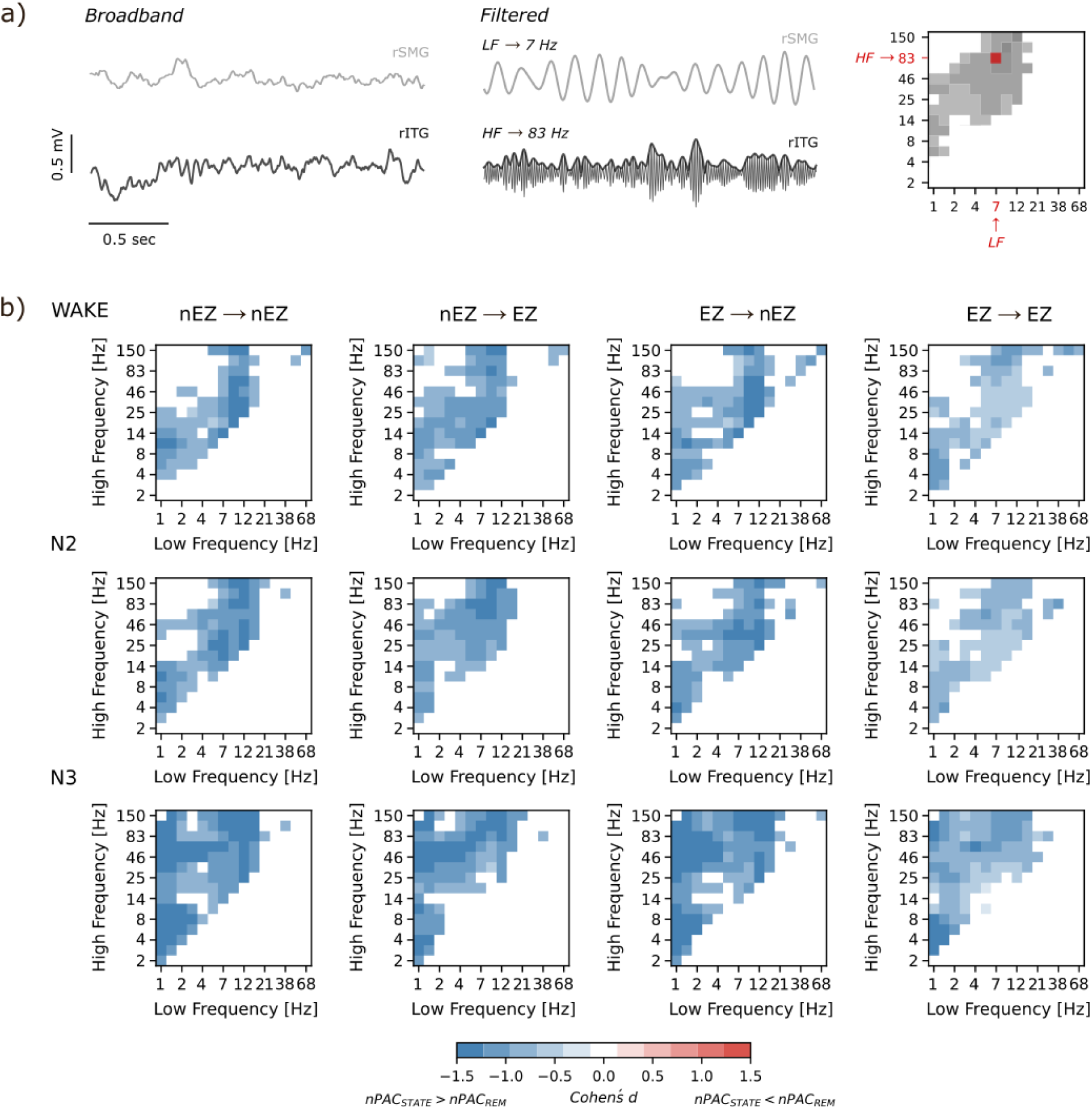
Reduced cross-frequency coupling during REM. a) Phase-Amplitude Coupling (PAC) computation between a single channel pair (LF: 7 Hz, HF: 83 Hz; right inferior temporal gyrus (rIGT) and right supramarginal gyrus (rSMG)); b) Panels display effect sizes of the PACs differences across states for: nEZ → nEZ, nEZ → EZ, EZ → nEZ and EZ → EZ. Effect sizes are represented as Cohen’s d values and only significant values (*p* < 0.05, Wilcoxon signed-rank test, BH-corrected) are displayed.

Compared to N3, REM sleep exhibited the largest differences (Fig. 3b), with a marked reduction (*p* < 0.05, d < −0.8) of the established slow-oscillations-driven modulation characteristic of N3, reaching a maximum decrease of approximately 30%. This attenuation involved not only δ-phase driven coupling but also the coupling between α–low-β phases and high-β–γ amplitudes. A similar pattern of reduction, affecting specifically δ-phase to α-amplitude and α–low-β to high-β–γ interactions, was observed when comparing REM to the other vigilance states across all channel-pair configurations. However, the magnitude of the effect was smaller in EZ → EZ connections (Fig. 3b).

These findings suggest that REM sleep is characterized by a marked decrease in cross-frequency interactions.

### REM sleep is associated with lower bistability across frequencies

We previously reported that larger β–γ band bistability indicates increased instability of oscillations and it has been associated with elevated seizure risk during both resting-state and NREM sleep SEEG recordings. We next examined whether REM sleep differs from other vigilance states in neuronal oscillation bistability (BiS) in both nEZ and EZ regions..

REM showed a wide-band large reduction of bistability compared to the other vigilance states (Fig. 4b, Suppl. Fig. 2), with both N2 and N3 showing significantly higher bistability compared to REM sleep, with a maximum of decrease 17% compared to N2 and 23% against N3, (*p* < 0.05, Wilcoxon signed-rank test, BH-corrected, Cohen’s d > 0.8; Fig. 4b, Suppl. Fig. 2). Compared to wake it showed a smaller effect and narrow-band reduction.

**Fig 4:**
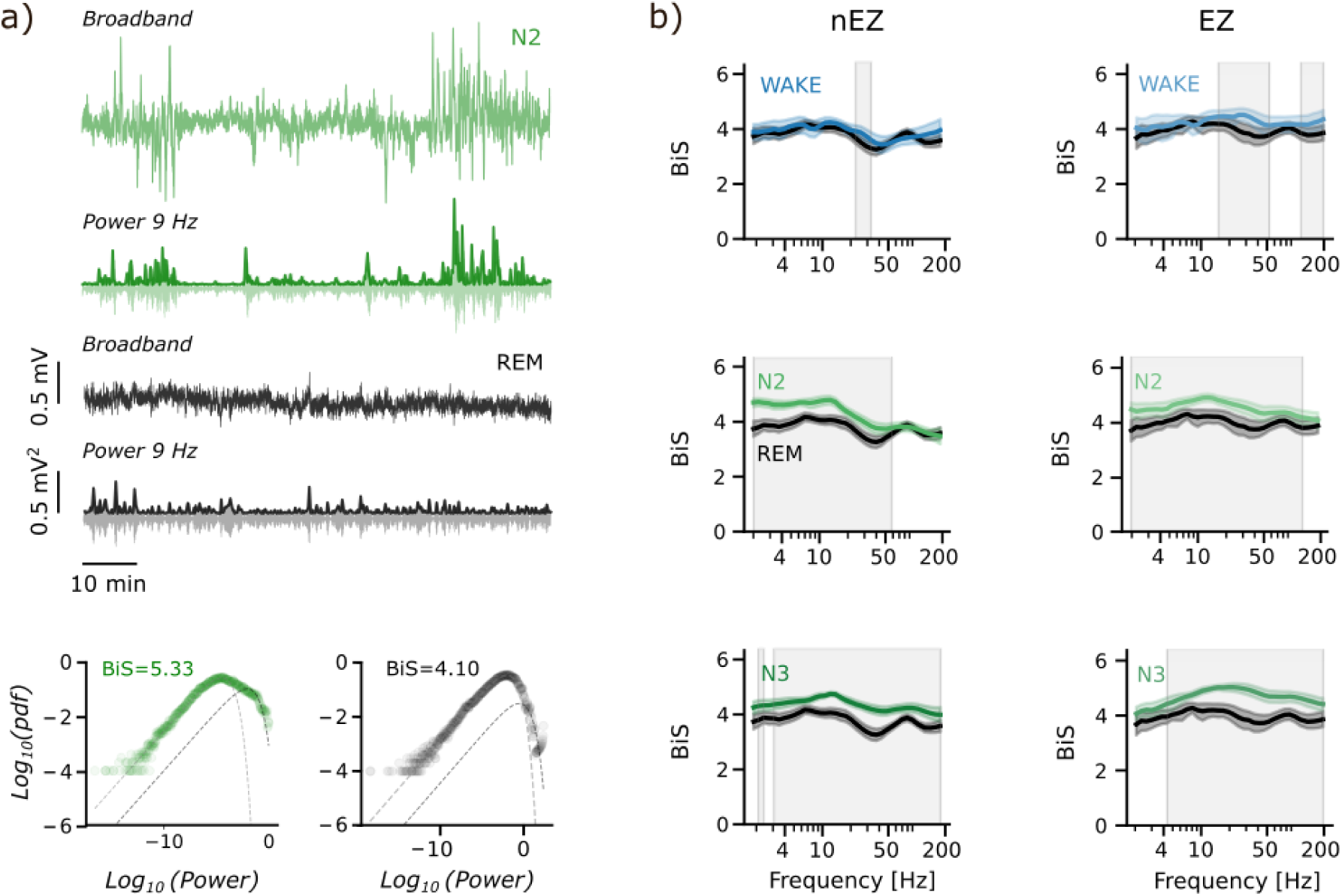
REM sleep is associated with lower bistability across frequencies. a) Individual-level evidence of differences in broadband signals and power time-series between REM and N2 sleep stages, with the corresponding fitting of the BiS index for a single contact located in the right supramarginal gyrus of a representative subject; b) BiS spectrum for nEZ and EZ, with colored shaded areas representing bootstrapped, two-tailed 95% confidence intervals (N=1000); The grey vertical bands highlight frequency ranges with significant differences (*p* < 0.05, Wilcoxon signed-rank test, BH-corrected).

In nEZ, N3 was associated with a widespread increase in bistability (*p* < 0.05, Cohen’s d > 0.8), whereas N2 showed an increase in all the frequency ranges except for high-γ range (< 60 Hz, *p* < 0.05, Cohen’s d > 0.8) (Fig. 4b, Suppl. Fig. 3).

In EZ instead N3 exhibited a higher BiS compared to REM ranging from *θ* to low-γ (4–50 Hz, *p* < 0.05, Cohen’s d > 0.8), while N2 exhibited higher BiS across the entire spectrum (*p* < 0.05, Cohen’s d > 0.8) (Fig. 4b, Suppl. Fig. 3).

Between REM sleep and the resting state, we did not observe significant differences except for high β–γ bands (>20 Hz, Fig 4b, Suppl. Fig. 3), where wake showed a slightly higher BiS (3% increase for nEZ, 7% increase for EZ, *p* < 0.05), even though the effect was less evident (Cohen’s d = 0.5–0.8).

Overall, REM sleep was consistently associated with a broadband reduction in bistability when compared to all other sleep states.

### REM sleep shows consistently weaker canonical correlation between PAC, bistability, and synchrony

We recently proposed that enhanced epileptogenicity during N2 requires a tight interaction between δ-synchrony and β-γ bistability mediated δ-phase amplitude coupling of faster oscillations (Burlando et al., 2025). Here we aimed to assess whether this oscillatory coupling between phase of slow-oscillations to high-frequency amplitude bistability generalizes across vigilance states and we hypothesized that REM would present the lowest canonical correlation compared to the other vigilance states. Using canonical correlation analysis (CCA) across all vigilance states and along the four configurations, we found a significant, we found a significant (permutation test, N=10000, BH-corrected) lower correlation in REM (*ρ* ≈ 0.59 − 0.87) in all the channel configurations. In nEZ–nEZ, inward PAC–BiS coupling was numerically lower in REM than N2 (not significant) and higher in REM than N3.

Altogether these results generalize previous observation on the canonical correlation between synchrony and bistability, further suggesting that during REM sleep this correlation is significantly reduced.

**Tab. 1:**
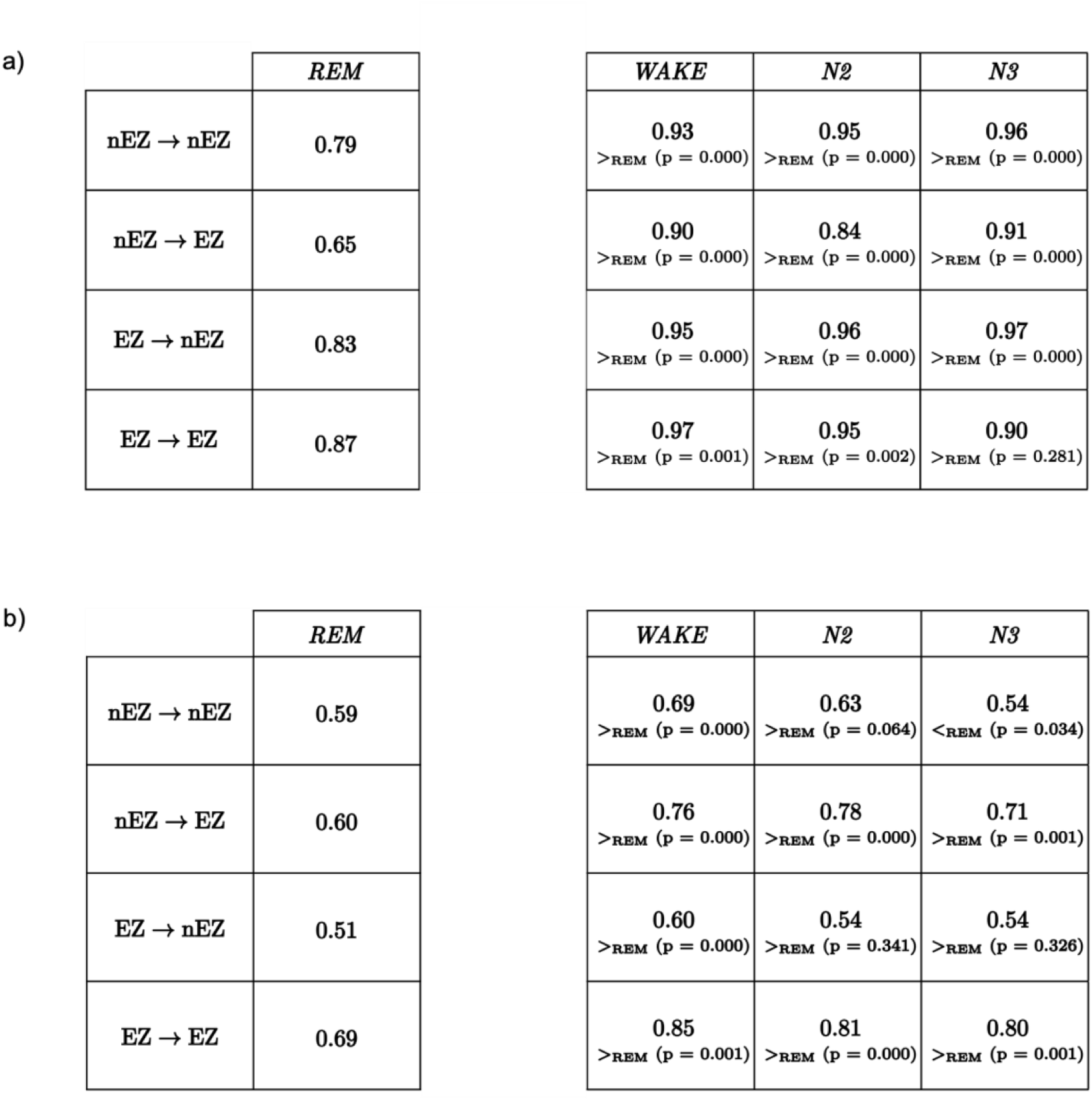
Canonical correlation coefficients comparison between REM and the others vigilance states across nEZ→nEZ, EZ→EZ, and nEZ→EZ. Table a) shows the canonical correlation coefficients obtained between phase-based metrics: SNS and Outward PAC in the 4 configurations. Table b) displays the canonical correlation coefficients computed between amplitude-based metrics: BiS and Inward PAC in the 4 configurations.

|  | *REM* |
| --- | --- |
| nEZ → nEZ | 0.79 |
| nEZ → EZ | 0.65 |
| EZ → nEZ | 0.83 |
| EZ → EZ | 0.87 |

| *WAKE* | *N2* | *N3* |
| --- | --- | --- |
| 0.93<br>><sub>REM</sub> (p = 0.000) | 0.95<br>><sub>REM</sub> (p = 0.000) | 0.96<br>><sub>REM</sub> (p = 0.000) |
| 0.90<br>><sub>REM</sub> (p = 0.000) | 0.84<br>><sub>REM</sub> (p = 0.000) | 0.91<br>><sub>REM</sub> (p = 0.000) |
| 0.95<br>><sub>REM</sub> (p = 0.000) | 0.96<br>><sub>REM</sub> (p = 0.000) | 0.97<br>><sub>REM</sub> (p = 0.000) |
| 0.97<br>><sub>REM</sub> (p = 0.001) | 0.95<br>><sub>REM</sub> (p = 0.002) | 0.90<br>><sub>REM</sub> (p = 0.281) |

b)
|  | *REM* |
| --- | --- |
| nEZ → nEZ | 0.59 |
| nEZ → EZ | 0.60 |
| EZ → nEZ | 0.51 |
| EZ → EZ | 0.69 |

**Tab. 1: Canonical correlation coefficients comparison between REM and the others vigilance states across nEZ→nEZ, EZ→EZ, and nEZ→EZ.**
| *WAKE* | *N2* | *N3* |
| --- | --- | --- |
| 0.69<br>><sub>REM</sub> (p = 0.000) | 0.63<br>><sub>REM</sub> (p = 0.064) | 0.54<br><<sub>REM</sub> (p = 0.034) |
| 0.76<br>><sub>REM</sub> (p = 0.000) | 0.78<br>><sub>REM</sub> (p = 0.000) | 0.71<br>><sub>REM</sub> (p = 0.001) |
| 0.60<br>><sub>REM</sub> (p = 0.000) | 0.54<br>><sub>REM</sub> (p = 0.341) | 0.54<br>><sub>REM</sub> (p = 0.326) |
| 0.85<br>><sub>REM</sub> (p = 0.001) | 0.81<br>><sub>REM</sub> (p = 0.000) | 0.80<br>><sub>REM</sub> (p = 0.001) |

## Discussion

In this study, we aimed to evaluate network-level dynamics to investigate on potential differences across vigilance states. We also wanted to evaluate whether REM shows a reduction of the pre-identified tripartite interaction linking neuronal synchronization, oscillatory amplitude bistability, and cross-frequency phase–amplitude coupling that may be linked to its suppressive effect on interictal discharges and seizures. We hypothesized that this protective role may be related to an attenuation of large-scale network processes that could otherwise promote hyper-synchronization and facilitate the propagation of epileptic activity.

Our results showed that REM sleep is characterized by a broadband reduction in neuronal synchrony, most evident in the θ–α range, together with weaker cross-frequency coupling and a significant decrease in bistability across a wide frequency spectrum. Previous work suggested that during NREM sleep, and particularly in N2, δ-synchrony promotes β–γ bursts through phase–amplitude coupling (PAC), thereby enhancing network instability and epileptogenic risk (Burlando et al., 2025). Our findings indicate that during REM, not only the δ-driven dynamic, but also a broader frequency-spanning coordination process (synchrony and cross-frequency coupling) is attenuated.

### REM suppresses large-scale synchronization

We found that REM sleep is characterized by a marked decrease in phase synchronization, focused in the θ–α range, across the three channel-pair configurations (EZ, nEZ, and nEZ–EZ, Fig. 1a) and stages.

In contrast, β–γ synchronization partly persisted during REM. This activity likely reflects physiological processes rather than pathological ones, as β and γ bands are involved in normal cognitive functions such as working memory consolidation (Kumral et al., 2025) and emotional and cognitive processing during dreaming (Walker, 2009).

Primarily in nEZ and nEZ–EZ, we also observed higher δ-band synchrony in REM, consistent with previous findings that a 1.5–3 Hz rhythmic slow oscillation characterizes human REM sleep, originating from mesiotemporal regions and resembling the hippocampal θ of animals (Bódizs et al., 2001; Moroni et al., 2012, 2007). This reduction was predominantly observed in comparison to N3, in line with previous studies that have reported limited evidence for robust large-scale delta synchronization during N3 (Cox et al., 2020).

N3 sleep is dominated by cortical delta oscillations (De Andrés et al., 2011), prior evidence suggests that these oscillations can propagate as travelling waves across widespread cortical regions (Massimini et al., 2004). However, the detection of such phenomena may depend on the spatial scale of the recording technique. SEEG provides high spatial resolution but samples neural activity locally. In contrast, studies demonstrating delta travelling waves during N3 typically rely on recording techniques that allow the examination of extended cortical areas (Massimini et al., 2004). Therefore, the absence of a marked increase in δ-band phase synchronization during N3 in our data likely limited spatial sampling of SEEG recordings rather than a genuine lack of delta oscillations propagation.

### REM reduces cross-frequency coupling

Phase–amplitude coupling (PAC) links slow network rhythms to faster local activity and can support physiological communication across structures. During REM sleep, PAC is still present, δ phase coupling with γ amplitude has been reported as a process sustaining hippocampal–cortical interactions in this stage (Clemens et al., 2009), yet its overall magnitude is lower than in other stages (von Ellenrieder et al., 2020).

We observed a generalized decrease in nPAC during REM sleep, particularly between δ phases and α amplitudes, as well as between α – low-β phases and γ m amplitudes. The largest reduction was found in comparison with N3, consistent with previous findings (Amiri et al., 2016). Stronger PAC in epileptogenic regions has been associated with pathological neuronal synchrony, where abnormal coupling between slow and fast oscillations emerges from hyper-synchronous neuronal firing driven by the excitatory phase of slow rhythms (Amiri et al., 2016). Conversely, a reduction in PAC may reflect a preserved excitatory–inhibitory balance and the absence of pathological hyper-synchronization (Amiri et al., 2016). Within this framework, the global reduction of nPAC observed during REM sleep may reflect a state of reduced pathological synchronization that might mitigate the excessive coupling that arises between slow and fast oscillations typical of epileptic networks.

### REM is associated with lower bistability

We found that REM sleep was characterized by a broadband reduction in bistability compared to all other vigilance states, primarily relative to N2 and N3, which instead showed markedly higher BiS values across a wide frequency range consistent with previous report (Burlando et al., 2025; Wang et al., 2024, 2023). This attenuation was evident in both epileptogenic and non-epileptogenic regions. In the context of epilepsy, elevated bistability reflects an increased propensity of cortical circuits to switch between stable and hyper-excitable states, favoring the emergence of epileptiform discharges (Fuscà et al., 2023; Meisel et al., 2015; Nobili et al., 2025; Wang et al., 2024).

Therefore, the reduction in BiS observed during REM may reflect enhanced network stability, characterized by a diminished susceptibility to pathological state transitions that could reduce likelihood of interictal or ictal activity.

### REM weakens the coupling between epileptogenic dynamics

Previous evidence showed that during NREM sleep, δ oscillations have been shown to modulate β–γ activity through PAC, linking slow-wave instability to increased bistability and the emergence of pathological fast bursts (Burlando et al., 2025). Based on this, we hypothesized that REM sleep, characterized by a broader variety of oscillatory rhythms and reduced slow-wave dominance than NREM (Cantero et al., 2002; Clemens et al., 2009; Hutchison and Rathore, 2015; Vijayan et al., 2017), attenuates not only the δ→β–γ coupling but also broader frequency-spanning coordination (i.e., large-scale synchrony and cross-frequency coupling).

To test this, we extended our analyses beyond δ to β–γ interactions to the entire frequency spectrum. Canonical correlation analysis revealed that REM sleep was associated with markedly weaker correlations between phase synchrony (node strength from PLV) and outward PAC with the respect to all the other vigilance states, as well as between inward PAC and bistability, compared to N2 and wakefulness.

The reduction of these correlations indicates that REM sleep may not only suppress each process independently - phase synchrony, cross-frequency coupling, and bistability - but also weaken their mutual relationships, thereby probably enhancing its role in limiting epileptiform activity.

Beyond this the lower correlation found between inward PAC and bistability in nEZ during N3 with the respect to other states supports the idea that N3 may not be strongly epileptogenic.

Altogether, these findings extend previous evidence of δ to β–γ oscillatory coupling during NREM sleep, showing that REM disrupts the coordination between phase- and amplitude-based processes, thereby limiting epileptogenic network dynamics, whereas NREM confirms its pro-epileptogenic nature by exhibiting the strongest correlations relative to REM.

### Conclusions and limitations

The suppressive effect observed during REM sleep likely arises from the combined influence of large-scale network desynchronization and neurochemical properties specific to this stage. REM is characterized by enhanced cholinergic tone, which promotes asynchronous neuronal firing and reduces the spatio-temporal summation of excitatory inputs (Clemens et al., 2009; Nir and Tononi, 2010). The co-occurring withdrawal of orexinergic activity, which normally supports cortical synchrony, further contributes to this desynchronized network configuration (Berteotti et al., 2023; Ng, 2017; Roliz and Kothare, 2022). Another signature feature of REM sleep is the occurrence of ponto-geniculo-occipital (PGO) waves, transient phasic events that transiently increase neuronal firing but have been associated with reduced seizure frequency, shorter seizure duration, and prolonged seizure latency (Salado et al., 2008). Together, these features provide a plausible physiological basis for the evidenced suppression of epileptogenic activity observed during this stage.

In line with these mechanisms, our findings demonstrated that REM sleep was associated with a marked attenuation of large-scale epileptogenic dynamics as well as a weakening of their mutual canonical correlations. This broad decoupling may reflect a shift toward a more physiological neuronal communication, reducing the likelihood of pathological state transitions and the propagation of epileptiform activity across cortical networks. This reduction was observed primarily relative to NREM sleep and, notably, also relative to resting wakefulness, which shares similar EEG properties with REM (Blumberg et al., 2020; Scammell et al., 2017).

In our study, epileptic discharges were not directly quantified, however previous investigations consistently reported their decrease during REM sleep (Campana et al., 2017; Frauscher et al., 2016). The next step could be to directly examine the relationship between the occurrence of epileptic discharges and the large-scale network dynamics we’ve evaluated here.

Moreover, prior evidence indicates that the phasic phase of REM, characterized by bursts of rapid eye movements and enhanced cholinergic drive with more marked cortical desynchronization, exerts an even stronger inhibitory influence on epileptic activity than tonic REM (Campana et al., 2017; Frauscher et al., 2016; Nobili et al., 2025). Recent studies have investigated on the network dynamics differences between these two substates of REM (Avigdor et al., 2025; Campana et al., 2017; Simor et al., 2021, 2020). Future research could then aim to better clarify how tonic and phasic REM differentially modulate these large-scale dynamics in the context of epilepsy.

We acknowledge some limitations, starting from the relatively limited sample size that may reduce statistical power. In addition, we couldn’t control for clinical factors such as pharmacological treatment, disease duration, lesions presence and the different types of etiopathology. Further investigation in larger, clinically stratified cohorts will be important to test the generalizability of these findings.

Notably, the suppressive effect of REM emerged consistently across patients with heterogeneous epilepsy types and independently of the anatomical localization of the epileptogenic zone, suggesting that these protective mechanisms might be independent from the specific epilepsy type and brain area.

Although EZ labels were defined from presurgical SEEG patterns (including early propagation regions), postsurgical outcome confirmed correct localization in only 65% of operated patients. Thus, some contacts classified as EZ or nEZ may not have perfectly matched the true epileptogenic tissue. However, a suboptimal Engel outcome does not necessarily imply mislocalization, as it may reflect incomplete treatment of a distributed epileptogenic network, limited sampling, multifocality, or surgical constraints. Importantly, any misclassification would likely reduce the contrast between EZ and nEZ rather than artificially produce a REM–NREM separation. The consistent REM-related attenuation observed across synchrony, PAC, bistability, and their coupling therefore supports the robustness of our main conclusions on vigilance-state modulation.

Altogether, our findings align with previous evidence that REM sleep is a vigilance state in which large-scale synchrony and cross-frequency coupling diminish, and they extend this picture by revealing a broadband suppression of bistability across both EZ and nEZ tissue, with canonical correlation analysis showing that REM also weakens the coupling among pro-epileptogenic mechanisms themselves. This combined attenuation of the individual processes and of their interdependence emerges not only relative to NREM but also to resting wakefulness, showing that REM likely reduces the processes that sustain hyperexcitability and the propagation of epileptiform activity.

