## Supplementary Material for "REM sleep reconfigures large-scale network dynamics: a link to its suppressive role in epilepsy"

### Supplementary, Figure 1

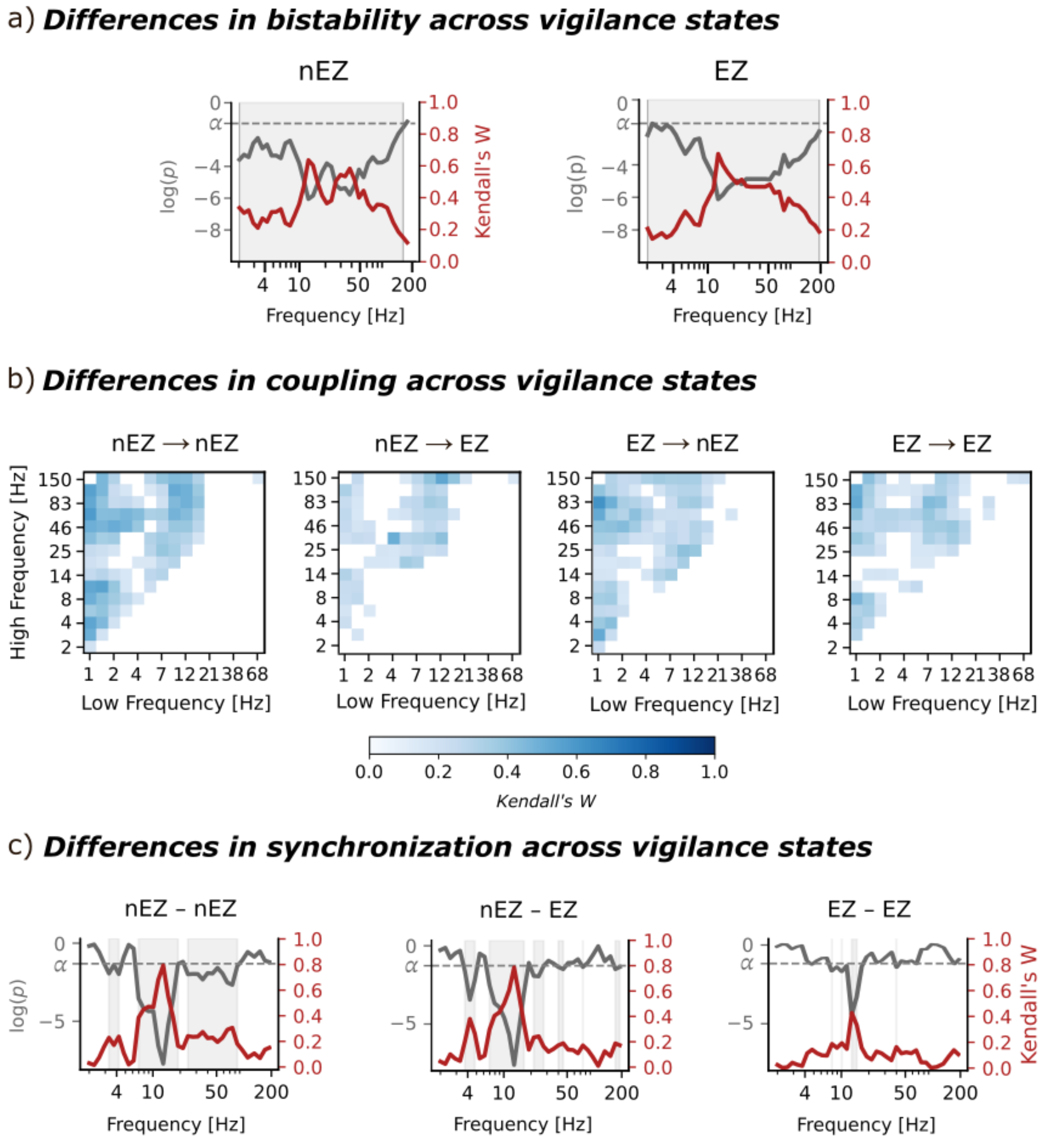


**PLV, nPAC, and BiS comparisons across vigilance-state and configurations**

Each row represents the comparisons respectively for a) PLV b) nPAC c) BiS.

In a) and c) p-values (Friedman test, BH-corrected) and effect sizes (Kendall's W) are shown across frequencies. The grey vertical bands highlight frequency ranges with significant differences ($p < 0.05$); In b) panels represent effect sizes of the PACs differences across states and configurations. Effect sizes are represented as Kendall's W values and only significant values ($p < 0.05$, Friedman test, BH-corrected) are displayed.

### Supplementary, Figure 2

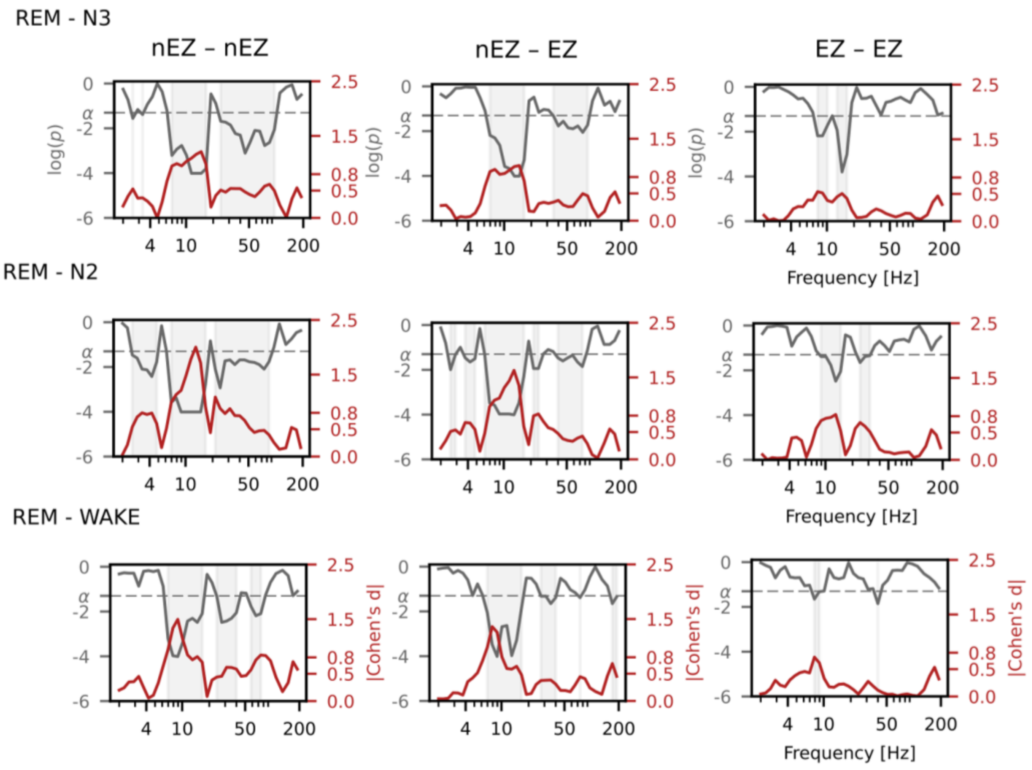


**PLV paired test between REM and the other vigilance states for nEZ-nEZ, nEZ-EZ, EZ-EZ.** Each panel represent p-values (Wilcoxon signed-rank test, BH-corrected; grey lines) and effect sizes (|Cohen's d|; red lines), the grey dashed line represent the significance level ($\alpha$).
Grey vertical bands highlight frequency ranges with significant differences ($p$ < 0.05) in PLV.

### Supplementary, Figure 3

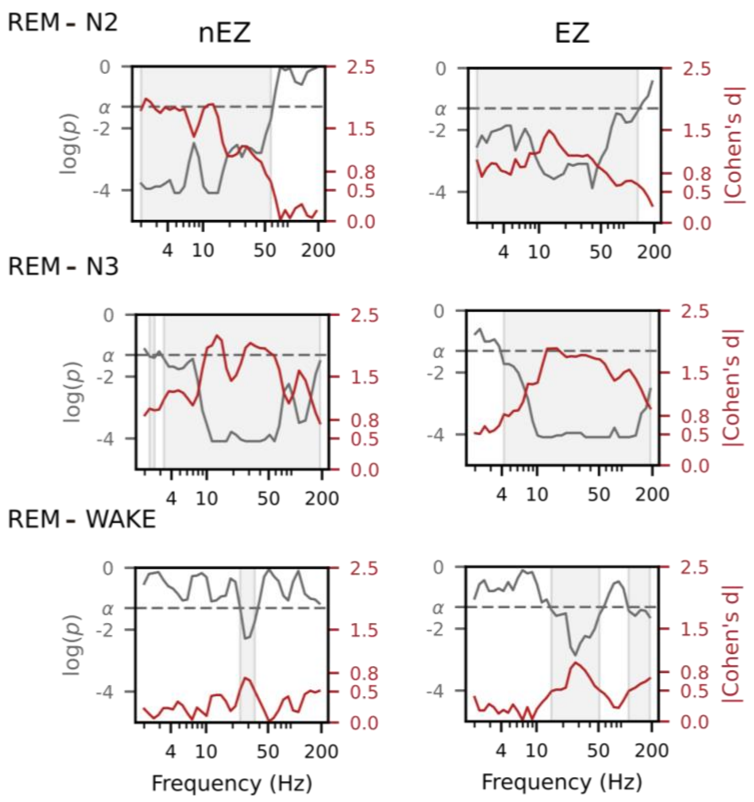


**BiS paired test between REM and the other vigilance states for nEZ and EZ.**
Each panel represent p-values (Wilcoxon signed-rank test, BH-corrected; grey lines) and effect sizes (|Cohen's d|; red lines), the grey dashed line represent the significance level ($\alpha$).
Grey vertical bands highlight frequency ranges with significant differences ($p$ < 0.05) in PLV.

### Supplementary, Figure 4

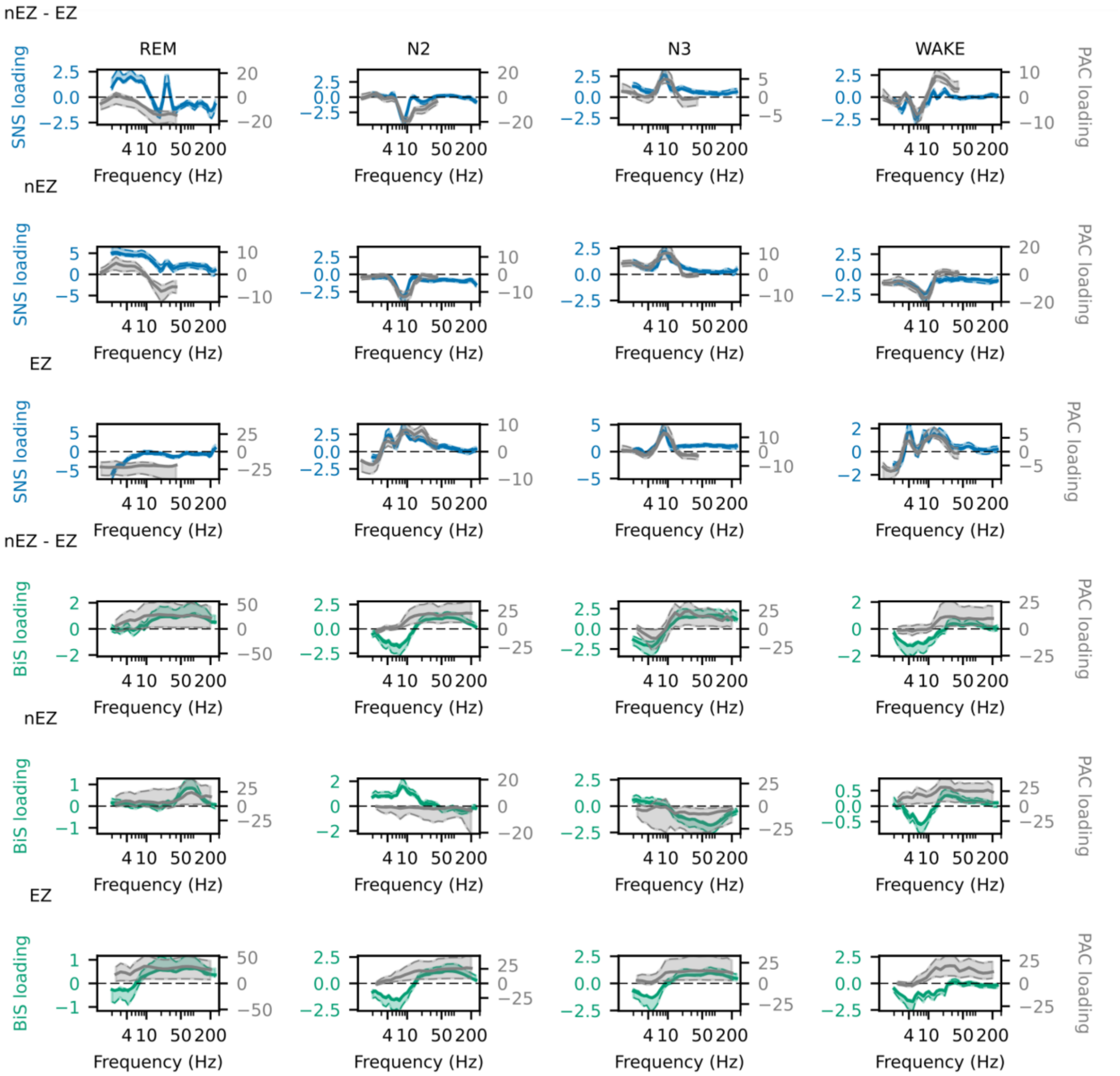


**Canonical correlation loadings between REM sleep with N2, N3, and Wakefulness** across nEZ→nEZ, EZ→EZ, and nEZ→EZ. The upper panels in each row display the CCA loadings spectra of Outward PAC (grey line) and SNS (green line), while lowers panels shows the CCA loadings spectra for the Inward PAC (grey line) and BiS (blue line). Colored shaded areas representing 95% confidence intervals (2.5%–97.5%).
